# Tailor: non-parametric and rapid score calibration method for database search-based peptide identification in shotgun proteomics

**DOI:** 10.1101/831776

**Authors:** Pavel Sulimov, Attila Kertész-Farkas

## Abstract

Peptide-spectrum-match (PSM) scores used in database searching are calibrated to spectrum- or spectrum-peptide-specific null distributions. Some calibration methods rely on specific assumptions and use analytical models (e.g. binomial distributions), whereas other methods utilize exact empirical null distributions. The former may be inaccurate because of unjustified assumptions, while the latter are accurate, albeit computationally exhaustive. Here, we introduce a novel, non-parametric, heuristic PSM score calibration method, called Tailor, which calibrates PSM scores by dividing it with the top 100-quantile of the empirical, spectrum-specific null distributions (i.e. the score with an associated p-value of 0.01 at the tail, hence the name) observed during database searching. Tailor does not require any optimization steps or long calculations; it does not rely on any assumptions on the form of the score distribution, it works with any score functions with high- and low-resolution information. In our benchmark, we re-calibrated the match scores of XCorr from Crux, HyperScore scores from X!Tandem, and the p-values from OMSSA with Tailor method, and obtained more spectrum annotation than with raw scores at any false discovery rate level. Moreover, Tailor provided slightly more annotations than E-values of X!Tandem and OMSSA and approached the performance of the computationally exhaustive exact p-value method for XCorr on spectrum datasets containing low-resolution fragmentation information (MS2) around 20-150 times faster. On high-resolution MS2 datasets, the Tailor method with XCorr achieved state-of-the-art performance, and produced more annotations than the well-calibrated Res-ev score around 50-80 times faster.

**Graphical TOC Entry:** 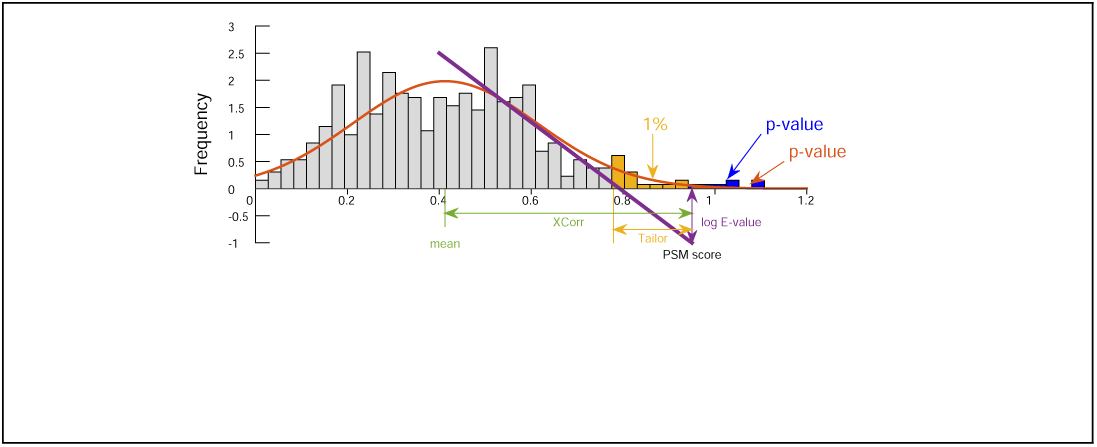

## Introduction

Mass spectrometry is now the de facto method used for protein identification in complex biological samples 1. Subsequent computational analysis of data involves the application of long pipelines of various algorithms to reveal the protein identification and structure^2–4^. Score functions are the workhorses in these pipelines. Each experimental spectrum obtained from an experiment is iteratively scored against a set of so-called candidate peptides, which are peptides that selected from a peptide database and whose calculated masses are equal to the precursor mass of the observed spectrum up to a certain tolerance. The scoring typically provides a similarity-like score based on matching the peaks of the experimental and the theoretical spectrum generated from the peptide sequences in silico. The top-scoring peptide-spectrum-match (PSM) is assigned to the experimental spectrum, and this assignment may be correct or incorrect. Uncalibrated, raw scores may indicate different match quality for different spectra. For instance, the distributions of the top scoring PSMs of doubly and triply charged spectra shown in Figure 1A indicate that a raw score of 2.5 may imply a correct annotation for a doubly but an incorrect annotation for a triply charged peptide molecule^5^. Spectrum-specific score calibration methods aim to provide a sort of score normalization so that spectrum assignments become comparable with each other; therefore, a single threshold can be selected to accept or reject spectrum annotations. The calibration allows one to obtain many more spectrum annotations at any desired false discovery rate (FDR)^5^. Score calibration methods involve a null distribution and calibrate a raw score to either the mean or the tail of the null distribution. In this article, we focus on spectrum specific score calibration methods and we disregard methods aimed at (a) assessing the confidence of the correctness of PSMs at whole-experiment level, such as q-values^6^, (b) improving protein identification^7,8^, (c) improving false discovery rate (FDR) estimation^9^, or (d) search protocols^10^.

**Figure 1:**
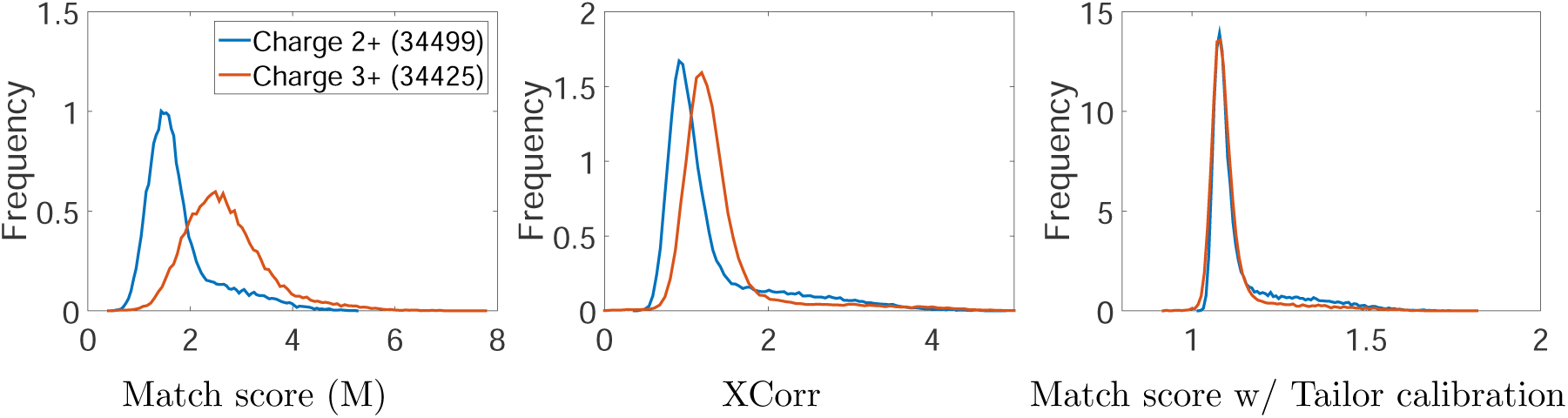
Distributions of top scoring PSMs obtained on Yeast data for doubly (blue) and triply (red) charged precursor ions. The panels show the distribution of the scores obtained with simple match score M(.,.) (A), with XCorr (B), and with match scores calibrated without the Tailor method (C). The density functions were obtained by kernel smoothing using the Gaussian smoother and normalized so that the area under the curves is equal to 1.0.

Every experimental spectrum can be associated with a null distribution, which may depend on the experimental spectrum itself, its precursor mass and charge state, the score function, the precision of the discretization, and on the theoretical peptide generation procedure (i.e. the charge states and the types, *b, y, a* etc., of the theoretical fragmentation ions). For instance, the XCorr^11^ score function implemented in Sequest 12,13, in Comet^14^ (an open source reimplementation of SEQUEST^12^), and in tide-search^15^ generates *y*^2+^, *b*^2+^ double charged theoretical fragmentation ion masses if the charge state of the precursor ion is higher than 2. Therefore, theoretical spectra contain twice as many peaks resulting in higher scores simply by chance. Another example is the score function of the Andromeda program^16^, which additionally generates water or ammonia loss peaks for peptides containing D,E,S,T or K,N,Q,R amino acids, respectively. Therefore, a single experimental spectrum can be associated with multiple null distributions with respect to the amino acid compositions of the theoretical peptides.

The XCorr score function is one of the most popular score function. It was developed for the SEQUEST ms/ms database search tool^11–13^ and it calibrates the match scores to the mean of the null distribution. The XCorr is defined as

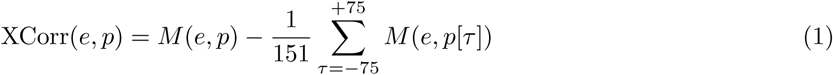

for a discretized experimental spectrum *e* and a theoretical spectrum *p* and consists of two parts. The first part simply qualifies the match between the experimental and theoretical spectra using the inner product of the corresponding vectors denoted by *M* (., .). The second part provides an estimation of the mean of the null distribution from 151 random matches obtained with a random theoretical peptide *p*[*τ*] generated by shifting the components of vector *p* by *τ* steps. We note that theoretical spectra corresponds to a real peptide sequence while shifting its components by ±*τ* > 0 steps breaks its semantics, and it cannot be associated with any real peptide sequence of the original mass, hence resulting in a random vector. Consequently, the XCorr score returns the signed difference between the match score and an estimated mean of the null distribution. Hence, XCorr incorporates a score calibration step, which is somewhat similar to the standardization (or Z-score normalization) defined as *z* = (*x* − *µ*)*/σ*, where *x, µ*, and *σ* represents the match score, and the mean and variance of the null distribution, respectively. We note that we tested a variation of the XCorr, in which the scores were normalized with the variance, but unfortunately, the variance can often be close to zero, and division by tiny numbers made the overall results worse. Figure 1B shows the distribution of the XCorr scores of the spectrum annotations separately for the doubly and triply charged spectra, and the figure suggests that the calibration mitigates the problem, albeit not entirely. We argue that the XCorr is suboptimal because it does not take into account neither the variance nor the tail of the null distribution.

The standard approach of score calibration is to assign a spectrum-specific statistical significance to a raw PSM score by estimating a probability of observing a random score equal to or greater than the observed PSM score. This is the p-value, which in fact has well-defined and accurate semantics^17^ over various experimental protocols and diverse configurations of MS instruments. The success of the score calibration methods relies on how well they approximate the tail or the extreme tail of the null distribution to obtain a p-value estimation. Some methods employ analytical models, such as a binomial distribution in Andromeda^16^ and MS Amanda^18^, Poisson distribution by OMSSA^19^,a Weilbul distribution for the XCorr^20^, or a Gumbel distribution for SSPV^21^, and rely on the assumption that peaks match independently between spectra. The disadvantages of these models include that (a) this assumption is not justified in practice^22^ and (b) the analytical probability mass functions (PMF) of binomial or Poisson distributions do not have cumulative distribution functions in closed forms to calculate the p-value instantly. As a result, they require a longer CPU time to sum over a larger number of PFMs at hypothetical PSM scores. The parameters of the exponential distributions (Weibull and Gumbel) are fitted from empirical PSM scores separately for each spectrum.

X!Tandem^23^ and Comet^14^ fits a linear regression line to the estimated survival function of the null model to calibrate the score for each experimental spectrum. Comet employs a log transformation of the survival function, and fits a linear regression line, and calculates a calibrated score, an E-value, by extrapolating the linear regression model at the top-scoring PSM score. X!Tandem employs a similar approach; it fits a linear regression line to the empirical survival function of the log of the HyperScores^23^. Both approaches assume that the tails of the null distribution decays exponentially; however, this assumption has not been critically analyzed.

The drawbacks of score calibration methods based on fitting specific parametric models includes that they cannot be straightforwardly generalized to other score functions and that the parametric distribution whose parameters are estimated using the overall distributions of PSM scores might not be accurate at the extreme tail^21^.

Other types of p-value estimation methods exploit the exact null distribution obtained from scoring all possible peptide sequences that have the same precursor mass as the observed spectrum^24–28^. The explicit enumeration of all sequences is computationally unfeasible; therefore, dynamic programming technique is employed to count the peptides at each score in the null distribution. These methods, indeed, result in a perfect score calibration; however, they have several drawbacks. First, they require proper estimation of the amino acid frequencies in the peptide database. Second, the calculation of the elements of the dynamic programming table requires a significant amount of CPU time. Third, the dynamic programming approach requires the score function to be additive^27^. Fourth, the dynamic programming method fails for peak-matching-based score functions (e.g. XCorr) used with data of high-resolution fragment mass accuracy because the fragmentation ions are approximated by the sum of the discretized masses of the amino acids in the dynamic programming method, which in turn can be different from the discretized mass of the whole fragmentation ion in high-resolution settings. We note that, for low-resolution MS2 data, the information loss due to discretization hardly poses any problems in practice. This is discussed in details by Lin et al.^28^. To overcome many of the issues mentioned above, empirical p-values of PSMs can be estimated via scoring spectra against a large number, say 10K, of decoy peptide databases^5^. In this scenario, well-calibrated p-values can be obtained for any type of score function using with high- or low-resolution MS2 data, albeit at the expense of CPU time. Figure 2 illustrates and compares the principles of the score calibration methods on a null distribution.

**Figure 2:**
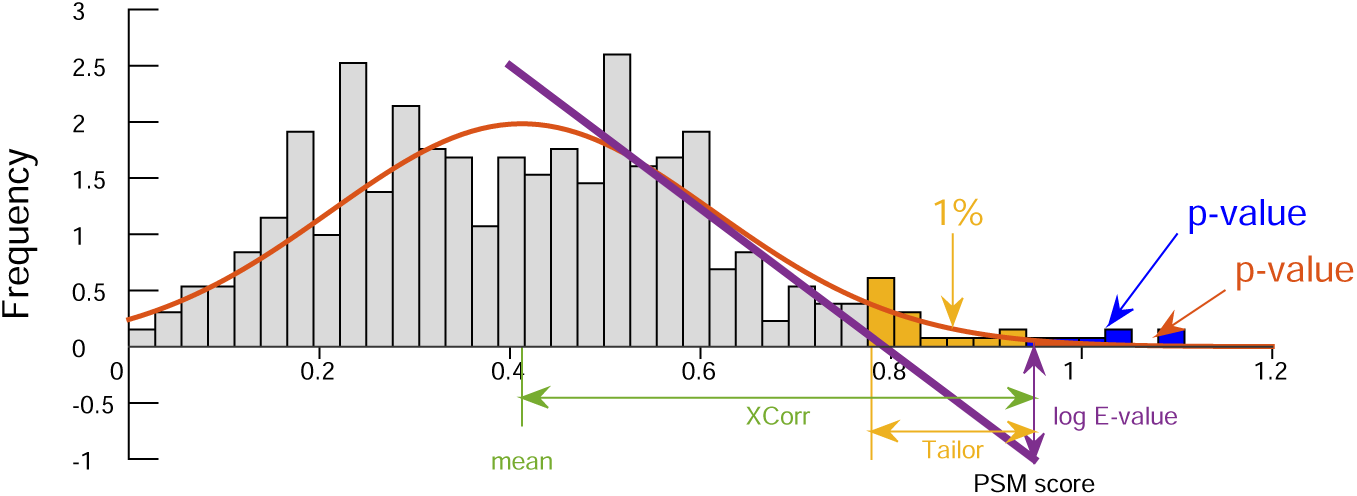
Illustration of the principles of the PSM score calibration approaches on a null distribution denoted by gray. (Green) XCorr calibrates the matching score by measuring the difference between the PSM score and an approximation of the mean of the random matching scores (Comet, Sequest, Tide). (Purple) Regression based methods fit a liner line on the empirical survival function based on the histogram of the random scores and extrapolates an E-value where a PSM score falls on this regression line (Comet, X!Tandem). (Blue) empirical p-values are calculated from the exact null distribution obtained with dynamic programming methods (XCorr exact p-value in Tide, MS-GF+) or with Monte-Carlo techniques^5^. (Red) P-values are calculated by using analytical probability density functions (OMSSA, Andromeda, Morpheus, SSPV, Weibull calibration of XCorrs). (Yellow) Tailor methods calibrates the score to the top 100-quantiles, i.e. relatively to the score which has a p-value of 0.01 (introduced in this article).

In this article, we propose a new, heuristic score calibration method, called Tailor, which calibrates the score of a PSM to the last 100-quantile, *Q*100, at the tail of the empirical null distribution, which is constructed for a given experimental spectrum from the scores obtained during scoring it against the candidate peptides. The Tailor method exploits the tail of the observed null distribution, which may be inaccurate but random scores are observed during the database search step, but not the extreme tail, where samples are rare. This is in contrast to the exact p-value methods (XPV, MSGF+), which enumerate all random scores, including those at the extreme tail, at the expense of CPU time to obtain an exact and accurate empirical null distribution. Therefore, Tailor is quick and works with any score function, albeit less accurate (i.e. a heuristic approach), whereas exact methods are accurate, albeit slow and require specific score functions.

Tailor is very easy to implement even in low-level programming languages such as C or C++, and it does not require specific statistical or optimization toolboxes. Tailor method is non-parametric, and it does not require any specific form of the null distribution. The main benefit of Tailor is its simplicity yet providing nearly perfect score calibration. For instance, the distributions of the spectrum assignment scores for doubly and triply charged experimental precursor ions calibrated with Tailor are nearly identical, as shown in Figure 1C. We emphasize that PSM scores ought to be calibrated for every experimental spectrum separately; however, we use the score distributions with respect to charge states only for illustrational purposes, as a sort of sanity checks.

This article is organized as follows. In the next section, we give a detailed definition of the Tailor method and explain the intuition behind it. Section provides details on our experimental protocols and the datasets we used. In Section, we show that Tailor yields significantly more spectrum annotations compared to raw and calibrated scores, such as E-values obtained with X!Tandem and OMSSA. We also show that Tailor performs comparably to the computationally intensive exact methods in spectrum annotation, although Tailor runs 20-160 times faster on our benchmark datasets. Lastly, Section presents the conclusions of our study.

## Tailor method

Let us consider an experimental spectrum *e* that is matched to *N* different candidate peptide sequences during the database searching step resulting in the following positive PSM scores: *s*_1_, *s*_2_, …, *s*_*N*_ > 0. Let us assume, for now, that *N* is large enough and that these scores are sorted in decreasing order; thus, the experimental spectrum *e* is to be annotated with the peptide sequence that produces the score *s*_1_. These scores form the basis of an empirical null distribution for the spectrum *e*. The 100-quantiles define 99 cut points dividing the range of the probability distribution into 100, continuous intervals with equal probabilities. The last (99th) score of the 100-quantiles of the empirical null distribution, denoted by Q100, is obtained here by selecting the PSM score at the position *i*\* = [*N/*100], where [.] denotes the standard rounding operation. Therefore, *Q*100 = *s*_*i*\*_ and the Tailor method calibrates the raw match scores by

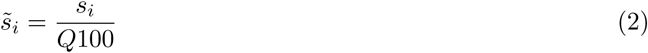

for *i* = 1, …, *N*, which are simply referred to as Tailor scores.

The Tailor score calibration is based on division instead of subtraction, and this relies on the following empirical observation. The mean of the null distribution is highly correlated with its standard deviation, as illustrated in Figure 3 for two experimental datasets from our benchmark. Consequently, a null distribution that has a larger mean also has a tail that decays slower compared to distributions that have a smaller mean. Therefore, a certain difference, say *l*, between the top score *s*_1_ and *Q*100 (*l* = *s*_1_ − *Q*100) might be significant for null distributions with a low mean but can be puny for those having a large mean. The ratio of *s*_1_ to *Q*100 implicitly takes into account the width of the null distribution; that is, the wider the null distribution, the higher its mean, the higher the score Q100; therefore, the score *s*_1_ is calibrated with a stronger factor *Q*100. Thus, the Tailor method incorporates the width (std) of the null distribution in this way. This is in contrast to the XCorr metric.

**Figure 3:**
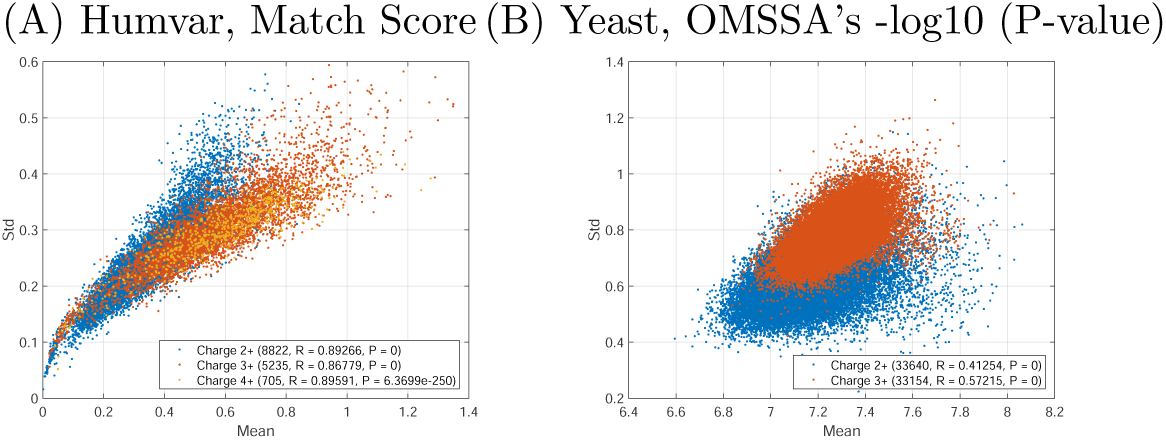
Correlation between the mean and the standard deviation (std) of the empirical null distributions. Each dot represents the mean and the std of the match scores of the candidate peptides of a single experimental spectrum. The scoring was carried out using the HumVar dataset with Match score (A) and Yeast with OMSSA’s -log(p-values) (B) against a decoy peptide dataset. For more information about the datasets used, see Section 3. Colors indicate charge states; the numbers in parentheses show the number of the spectra, the correlation coefficients (R), and the p-values (P) for testing the hypothesis that there is no relationship between the mean and the std (null hypothesis).

### Implementation details

The Tailor method is very simple to implement; we implemented it in the tide-search^15^ program in Crux^29^ using 13 lines of C++ code, and it is freely available in Crux at http:\\crux.ms. The Tailor method in tide-search can be activated using the parameter `--use-tailor-calibration T`, and the calibrated scores can be found in the column entitled `Tailor score`in the output.

We note that the number of candidate peptides *N* can be small, resulting in a degenerated null distribution. In this case, the Tailor method scores additional theoretical peptides that are the closest from the outside of the precursor mass window of a the experimental spectrum so that Q100 can be calculated based on at least 30 PSM scores (*N >* 30). We emphasize that, these PSMs are not taken into account in the peptide assignment. We also note that X!Tandem performs a similar approach, in which it generates additional peptide sequences with circular permutations in order to a build better null distribution for its E-value calculation. We also note that in a typical experiment, most spectra have more than 30 candidate peptides and this is a rather rare event. We also note that the position of Q100 is at least 3 (i.e. *Q*100 = *s*_*i*\*_, where *i*\* = min{3, [*N/*100]}) in order to avoid homologous or correlated peptides among the top-ranking PSMs^22^.

We also implemented the Tailor method in Python scripts to calibrate the score of the search results of X!Tandem and OMSSA. The Python scripts are provided in the supplementary materials. We note that in these scripts, Tailor eliminates any spectrum annotations that has fewer than 30 candidate peptides.

### Negative scores

Certain score functions (raw score from MSGF+, XCorr from Comet) may produce negative values for some PSMs, and it is very unfortunate when the signs of the score *s*_1_ and the Q100 are different. For such score functions, we add a constant value to every PSM scores large enough to keep all scores positive and larger than one.

## Datasets and methods

All data and scripts used in our benchmark are available in the Supplementary materials.

### Datasets

#### HumVar

The human variation dataset was derived from lymphoblastoid cell lines from 95 HapMap individuals, including 53 Caucasians, 33 Yorubans, 9 eastern Asians, and one Japanese^30^. The complete dataset is available at (http://www.peptideatlas.org/PASS/PASS00230), and it consists of 9,092,380 fragmentation spectra in 560 RAW files with a total size of 224 GB. In our study, we used only the `Linfeng_012511_HapMap39_3.mzML` file, which contained 15,057 experimental spectra. Protein lysates were subjected to detergent cleanup, cysteine alkylation, trypsin digestion, and isobaric tandem mass tag (TMT) labeling. Digested peptides were labeled with sixplex TMT, and the six TMT-labeled samples were equally mixed to generate the final digest mixture. All digest mixtures were analyzed on an LTQ Orbitrap Velos (Thermo Scientific) equipped with an online 2D nanoACQUITY UPLC System (Waters). During data acquisition, the full MS scan was performed in the orbitrap in the range of 400-1800 m/z at a resolution of 60000, followed by the selection of the 10 most intense ions for HCD-MS2 fragmentation using a precursor isolation window width of 1.5 m/z. Ions with a singly charged state or unassigned charge states were rejected for MS2. Ions within a 10 ppm m/z window around the ions selected for MS2 were excluded from further selection for fragmentation for 60 seconds. This dataset was searched against the IPI.Human database version 3.87, which contains 91,464 protein sequences. Peptides were generated using the trypsin cleavage rule, allowing a minimum length of 7 amino acids and up to one missed cleavages. Static carbamidomethylation (57.02146 Da) modification of cysteine, TMT labeling (229.16293 Da) of lysine and N-terminal amino acids, and oxidation (15.995 Da) on methionine were included as variable modifications. The database searching parameters for this dataset are summarized in Table 1.

**Table 1:** Summary of mass spectrometry datasets.

| Name | Instrument | #Spectra | Tolerance<sup>1</sup> | #Proteins<sup>2</sup> | #Peptides<sup>3</sup> | ACP<sup>4</sup> | MVM<sup>5</sup> | Modifications<sup>6</sup> |
| --- | --- | --- | --- | --- | --- | --- | --- | --- |
| HumVar | LTQ Orbitrap | 15,057 | 50 ppm / 0.02 Da | 91,464 | 3,420,673 | 1353.7 | 2 | O(V),TMT6-plex(V) |
| iPRG | MALDI 5600 | 14,141 | 10 ppm / 0.02 Da | 42,450 | 4,283,235 | 185.5 | 1 | O(V) |
| HSPP2A | LTQ | 29,583 | 50 ppm / 1.0005079 Da | 91,464 | 1,591,444 | 443.9 | 1 | O(V) |
| Yeast | LTQ | 69,705 | 3 Da / 1.0005079 Da | 6,734 | 269,373 | 702.4 | 0 | none |

<sup>1</sup>Precursor / fragmentation ion tolerance. No isotope error was allowed. <sup>2</sup>In silico enzymatic digestion was performed using the trypsin digestion rule allowing one missed cleavage. <sup>3</sup>Includes modified, target and decoy peptides. <sup>4</sup>Average number of candidate peptides per spectrum-charge combination (ACP). <sup>5</sup>Maximal variable modifications per peptide sequence. <sup>6</sup>Variable (V) and static (S) modifications, TMT-labeling (229.162932 Da) on lysine (K) and on N-terminal (Nt) modifications, oxidation (O) of methionine (+15.9949 Da). Static carbamidomethylation modification of cysteine (+57.02) was used for all datasets.

#### iPRG2012

The Proteome Informatics Research Group (iPRG) at the Association of Biomolecular Resource Facilities (ABRF) released an MS/MS dataset (in this paper called the iPRG dataset) to conduct a competition on detecting modified peptides in a complex mixture^31^. Further details and the evaluation of submissions from participants are available at ABRF’s website at https://abrf.org/research-group/proteome-informatics-research-group-iprg. The MS/MS dataset was obtained from tryptic digestion of yeast proteins and, in addition, was spiked with 69 synthetic modified peptides (called spike peptides). A candidate set of protein sequences (in fasta format) were also provided. Peptides were generated via the trypsin digestion rule. The static carbamidomethylation modification of cysteine (57.02146 Da) and one oxidation of methionine (15.9949 Da) per peptide were used as variable modification. The database searching parameters for this dataset are summarized in Table 1.

#### HSPP2A

This dataset was generated from the human protein phosphatase 2A system^32^ using an LTQ mass spectrometer, resulting in 29 583 spectra, and the data were downloaded from www.peptideatlas.org/repository/publications/Glatter2008 site. This dataset was searched against the IPI.Human database version 3.87, which contains 91,464 protein sequences. Peptides were generated using the trypsin cleavage rule, allowing a minimum length of 7 amino acids and up to one missed cleavages. Static carbamidomethylation (57.02146 Da) modification of cysteine and one variable oxidation (15.9949 Da) modification were included on methionine per peptide. The database searching parameters for this dataset are summarized in Table 1.

#### Yeast

This dataset was generated from yeast (*Saccharomyces cerevisiae* strain S288C) samples using a *µ*LC-MS/MS instrument, resulting in 69,705 spectra. This dataset was downloaded from https://noble.gs.washington.edu/proj/percolator/. Details on the sample preparation can be found in Section S1.3 of the supplementary materials of^33^. The yeast protein sequence file was also provided, containing 6734 protein sequences in fasta format. The peptide sequences were generated with trypsin cleavage using only the default carbamidomethylation modification of cysteine (+57.02146 Da) and allowing up to one missed cleavage. The database searching parameters for this dataset are summarized in Table 1.

### Database searching protocols and programs

The protein sequence (fasta) files were predigested in silico using the Pyteomics python toolbox^34,35^, allowing one missed cleavages. Decoy peptides were generated with the peptide-reverse approach, in which non-terminal amino acids were reversed. The peptides were exported in a fasta file format in which the labels “decoy” and “target” were appended to the header of the peptides. Therefore, database searching programs were not allowed to perform (a) any decoy peptide generation, (b) in silico protein digestion, and (c) false discovery rate (FDR) control. Each database searching program used the same peptide dataset with respect to the input dataset. The results were evaluated using the concatenated target-decoy approach^36^ on the output of the database search programs. The FDR was estimated as 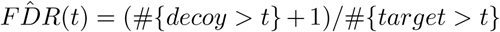, as proposed recently by^37,38^, and the q-values^39^ of the PSMs were calculated. The number of the accepted spectra were plotted at various q-value thresholds. In the resulting plots the higher curves stand for better performance.

In our benchmark, we used (a) the tide program from Crux v3.8.2, (b) X!Tandem v2015, and (c) OMSSA v2.1.9. The parameterization of these programs can be found in Supplementary Note 1. We emphasize that the Tailor method was implemented in the tide program, and the experiments were run under the same searching conditions (same search parameters, score function, filtering, preprocessing steps, peptide datasets, etc.). Consequently, the results obtained with different score calibration methods are directly comparable, and the gain or loss can be straightforwardly attributed to the score calibration methods, unless stated otherwise.

For X!Tandem and OMSSA, Tailor was implemented as a Python script and run on the output to recalibrate results. Therefore, the gain or loss in the search results within the same datasets can be directly compared and attributed to the score calibration methods.

We note that, the results obtained with the same datasets but using different database searching programs cannot be directly compared, because gain or loss in performance can arise from, for instance, different preprocessing steps or from the discriminative power of the different underlying score functions.

All programs were run on a Ubuntu Linux (v18.04.1) PC equipped with 64-bit Intel(R) Xeon(R) CPU E5-2640 v4 @ 2.40GHz, 128 Gb RAM. Scripts for plotting the results were written in Matlab.

## Results and discussion

### Robustness

We first show that the Tailor method is robust to parameterization. We executed the Tailor score calibration method using various quantiles as reference points on the Yeast dataset. This dataset was chosen because Yeast experimental spectra are associated with a large number of candidate peptides (702 candidate peptides per spectrum-charge combination on average) because of the low-resolution precursor information; thus, quantiles can be determined on large population. The following quantiles were studied: (a) the top 100-quantile denoted as Q100 and corresponding to a p-value of 0.01, (b) the top 20-quantile *Q*20 = *s*_min{3,[*N/*20]}_, (corresponding to a p-value of 0.05), (c) the top 5-quantile *Q*5 = *s*_min{3,[*N/*5]}_ (corresponding to a p-value of 0.2), and (d) the 2-quantile *Q*2 = *s*_min{3,[*N/*2]}_ (corresponding to a p-value of 0.5). The results are shown in Figure 4A and indicate that the quantile chosen as a reference point has little effect on the overall results, although there is a clear tendency. The closer the quantile is to the tail of the null distribution, the better the results are. The match scores calibrated to Q2 yields the lowest number of annotations, denoted by the purple line in this comparison; however, they still yield an improved spectrum annotation compared to the standard XCorr score function (marked with the green line). The match scores calibrated to Q100 yields the highest number of annotations, shown with the blue line, although Q20 closely follows (red line). Based on this experiment, we propose and use Q100 in the Tailor method. We note that the selection of the quantile does not have any effect on the run time.

**Figure 4:**
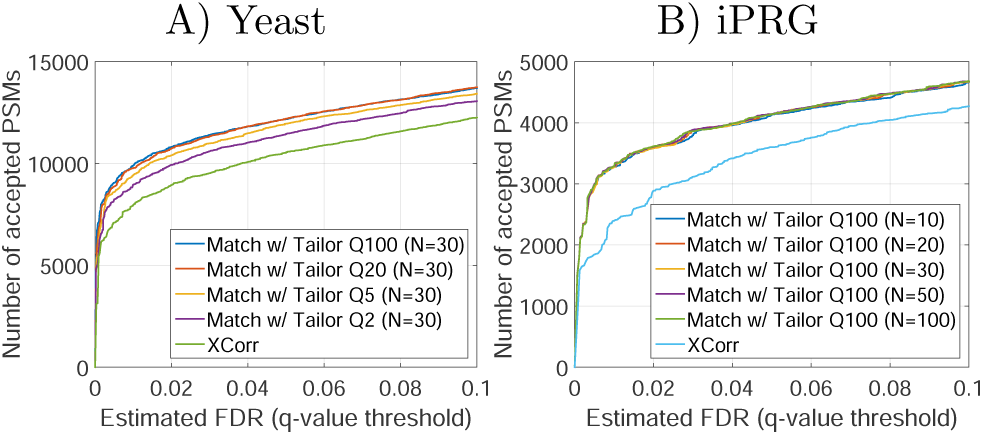
Robustness of the Tailor method to its parameter settings.

Next, we studied the effect of the threshold values (*N*) for the required minimal number of PSMs used to estimate the quantile score. This test was done using the iPRG dataset, because experimental spectra have relatively few candidate peptides because of the high-resolution precursor information; more specifically, there are 335, 572, 761, 1165, and 2337 experimental spectra, which have fewer than 10, 20, 30, 50, and 100 candidate peptides, respectively, out of a total 14,141 experimental spectra. The number of spectrum annotations shown in Figure 4B indicates that different thresholds result in a small variance in the outcome. Moreover, one could expect longer calculation time for higher threshold values; however, in practice, this has a negligible impact on CPU time. The run times were 18.3, 17.4, 19.4, 17.4, and 17.5 seconds for *N* =10, 20, 30, 50, and 100, respectively. We note that the run time for standard XCorr is 21.7 seconds. We conclude that the value of the threshold *N* has minimal impact on the search time and the results.

### Xcorr

The Tailor score calibration method was compared against the XCorr method^11^ and the XCorr’s exact p-value (XPV) methods^27^. The underlying score function of XCorr and XPV is the Match score. We obtained the results of the Match score by removing the `void SubtractBackground(double*, int)` procedure call (this procedure is responsible for performing the cross-correlation step) from the XCorr function in the `spectrum_preprocess2.cc` source file of tide-search. The numbers of the annotated spectra as a function of the FDR are shown in Figure 5, and the run times are shown in Table 2 separately for all spectrum datasets. The Match score, as expected, performs poorly on low-resolution MS2 data (HSPP2A, Yeast), as shown by the purple line, but when it is calibrated with the Tailor method, it consistently and significantly yields more peptides at any FDR than when it is calibrated with the cross-correlation method (i.e. the XCorr). These methods run very fast; however, the Tailor method is slightly faster than XCorr, because it does not require a cross-correlation step. The Tailor method approaches the exact and computationally expensive XPV score calibration method in spectrum annotation and even outperforms it on the Yeast dataset. The Tailor method is around 20 and 160 times faster than XPV on the HSPP2A and Yeast datasets, respectively.

**Table 2:** Database search time in seconds for various scoring and calibration methods.

| Name | HumVar | iPRG | HSPP2A | Yeast |
| --- | --- | --- | --- | --- |
| Match | 45.6 | 19.3 | 13.5 | 29.9 |
| Match w/Tailor | 48.4 | 19.4 | 13.6 | 30.4 |
| XCorr | 49.5 | 21.7 | 13.2 | 28.2 |
| XCorr w/ XPV | 318.0 | 138.0 | 309.0 | 4940.0 |
| Res-Ev<sup>1</sup> | 3400.0 | 308 | n.a. | n.a. |

<sup>1</sup>Res-ev was designed for high resolution MS2 data thus it was not run on the HSPP2A and Yeast datasets which contain low-resolution fragmentation ion information.

**Figure 5:**
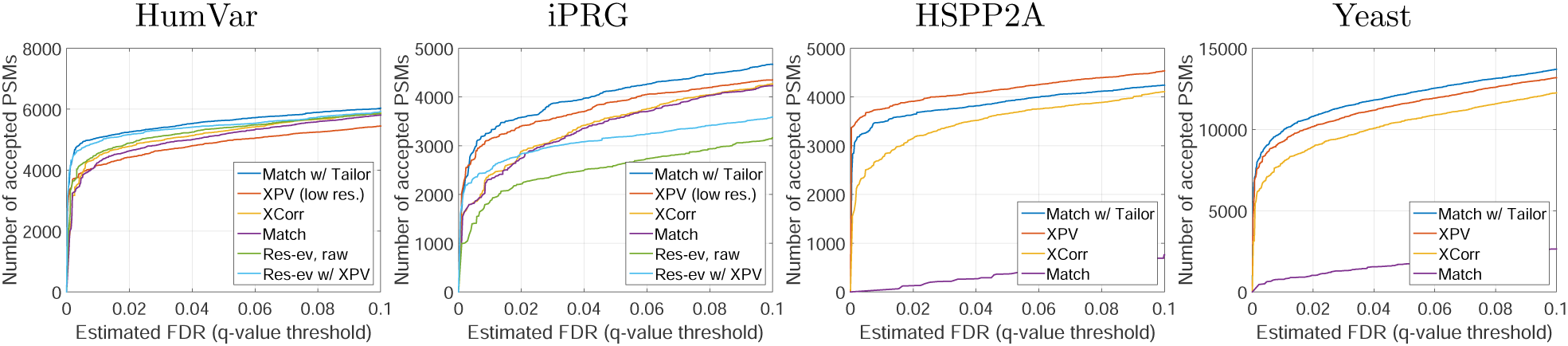
Results of spectrum annotation with various scoring and calibration methods using the tide program. Each plot corresponds to different datasets, as indicated on top. The top row corresponds to datasets of high-resolution fragmentation ion information, while the bottom row corresponds to datasets of low-resolution.

On high-resolution MS2 data (HumVar, iPRG), the raw Match score performs as well as the XCorr function, as shown by the purple and yellow lines in the top plots of Figure 5. This suggests that cross-correlation calibration for high-resolution MS2 data is futile, because, in our opinion, the mean of the null distribution (estimated by the cross-correlation part and then subtracted from the raw score) is close to zero. However, Tailor calibration can further increase the number of the annotated spectra at any FDR level significantly (compare the dark blue and yellow lines), because Tailor calibrates to the top percentile of the null distribution, which is possibly larger than zero.

We executed the XPV method as well, although with a wide bin `(bin_width=1.0005079)`because a narrow bin `(bin_width=0.02)` would break down XPV^28^. The results show that the Match scores that were corrected by the heuristic Tailor method executed with high-resolution settings yielded more spectrum annotations at any FDR level than the exact XPV methods with low-resolution settings in around 5-20 times less time. The residue-evidence (Res-ev) score function was design specifically for high-resolution MS2 data to overcome the shortcomings of XPV^28^ and it can be calibrated with exact methods using a dynamic programming approach. The calibrated Res-ev score (p-value) function yielded slightly fewer annotations on the HumVar dataset, and significantly fewer annotations on the iPRG dataset, at any FDR level than the Tailor method; compare the light and dark blue lines in the top plots of Figure 5. Furthermore, Tailor runs around 20-70 times faster depending on the dataset. We note that the results obtained with Match score with Tailor on high-resolution MS2 data cannot be directly compared to the results obtained with XPV using low-resolution MS2 data or to the Res-ev score using high-resolution MS2 data, because the gain or loss in spectrum annotation might not arise from the score calibration but from the discriminative capability of different score functions (i.e. the ability of separating correct annotations from incorrect ones^17^). However, these experiments show that Match score with Tailor correction can achieve state-of-the-art results on high resolution MS2 data. Moreover, the fact that the heuristic Tailor method outperformed the well-calibrated Res-ev method may suggests that the Res-ev score might be less discriminative than the XCorr, so there still is a room for improvement on datasets of high resolution.

Next, we plotted the score distributions of the top-scoring PSMs for all experimental spectra with respect to its charge state. The distributions are shown in the top row for the XCorr method and in the bottom row for the Tailor method in Supplementary Figure S1 for all datasets. The plots show that the modes of the XCorr score distributions have different locations with respect to the charge states. This is particularly well visible for the HumVar and Yeast datasets. Furthermore, the location of the modes vary over the datasets as well. The Figure 6A illustrates this for the distributions of the top-scoring PSMs of triply charged experimental spectra. The mode obtained for the HumVar datasets is located around 1.8, while the mode corresponding to the iPRG dataset is located around 0.5. The Tailor method does a better job on score calibration with respect to charge states shown in the bottom row of Supplementary Figure S1 and across different datasets shown in Figure 6B.

**Figure 6:**
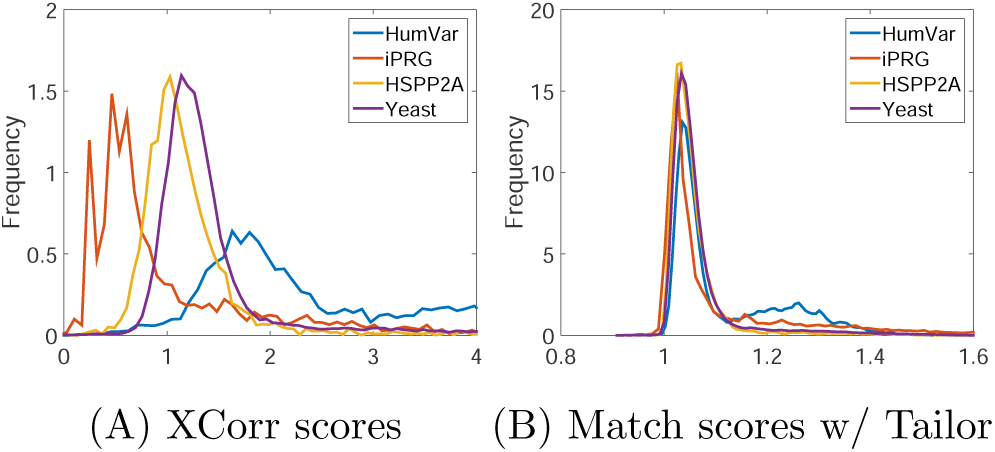
Distributions of the XCorr (LHS) and the Tailor (RHS) scores obtained using only experimental spectra that have an associated triply charged precursor ion. The distributions were smoothed using a Gaussian kernel.

### HyperScore

The Tailor score calibration was applied to the HyperScore metric of the X!Tandem search engine. The number of annotated experimental spectra obtained with raw HyperScore scores, E-values, and HyperScore scores calibrated with the Tailor method are shown in Figure 7. The results show that the Tailor method yields an improved number of spectrum annotations compared to the raw HyperScore, but Tailor is on par with the E-value method but yields more spectrum annotations than E-value only for the Yeast dataset. The score distributions smoothed with kernel smoothing with respect to charge states are shown in the top, middle, and bottom row of Supplementary Figure S2 for the raw HyperScore, the E-value, and the Tailor scores, respectively. These plots show that raw HyperScore results in different modes with respect to the charge states, whereas the E-values and the Tailor method results in modes roughly at the same location. The plots in panels A, B, and C of Figure 8 show the score distributions of the triply charged experimental spectra with respect to the dataset for the raw HyperScore, the E-value, and the HyperScore with Tailor method, respectively. These plots suggest that the E-values of the spectrum annotations and the Tailor calibration result in better score calibration compared to the raw HyperScore metric.

**Figure 7:**
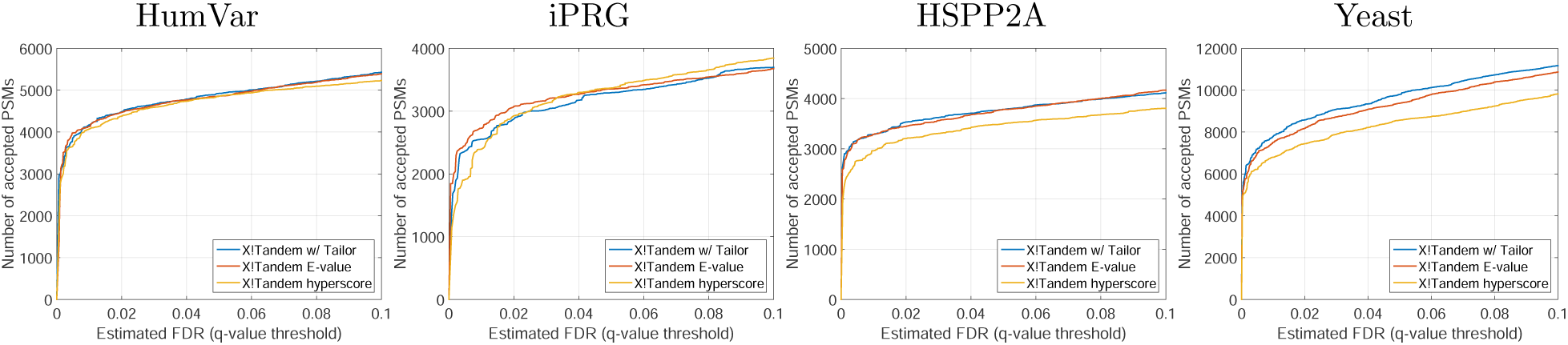
Spectrum annotation results using X!Tandem program. Each plot corresponds to different dataset indicated on top.

**Figure 8:**
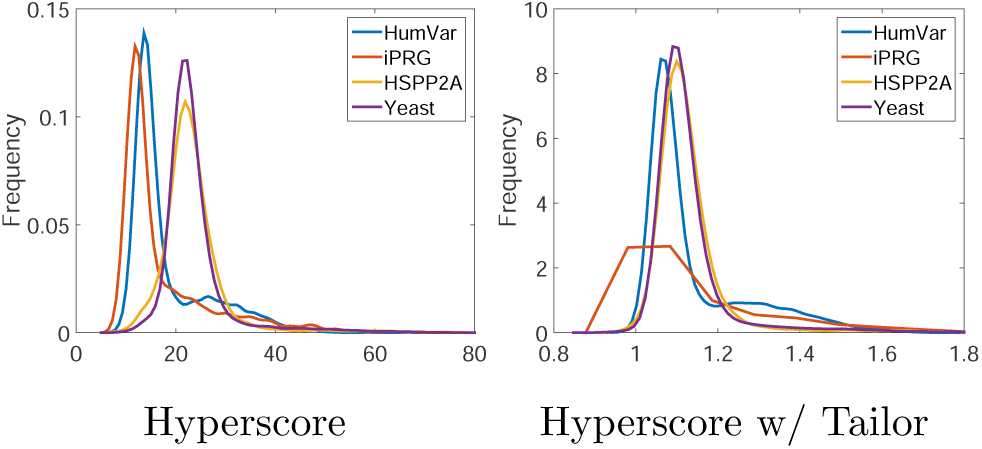
Distributions of the HyperScore (LHS) and Tailor (RHS) scores obtained using only experimental spectra having an associated triply charged precursor ion. Distributions were smoothed using Gaussian kernel.

### OMSSA score

We examined the p-values obtained with the OMSSA software and re-calibrated them with the Tailor method. We note that OMSSA does not report any raw match scores but only p-values and E-values of the corresponding PSMs, and the Tailor method was run to re-calibrate the minus-log transformation of the OMSSA’s p-values (i.e. − log10(*p* − *value*)). The transformation does not affect the FDR calculation but converts the p-values to a higher-the-better-like scores for Tailor and enables more appealing and informative visualizations.

The results plotted in Figure 9 show that the Tailor method managed to yield additional spectrum annotations by re-calibrating OMSSA’s p-values. This outcome suggests that the E-values could be further improved and OMSSA’s assumptions regarding its p-value calculation methodology might not be profoundly justified in practice. Supplementary Figure S3 plots the distribution of the minus-log transformation of the p-values (top row) and their calibration with the Tailor method (bottom row) with respect to the charge states. These plots indicate that the modes of these distributions are located around the some place for E-value and Tailor, except for the Yeast datasets, for which both methods produced different modes for doubly and triply charged spectra. However, we assume that the Tailor method “inherited” the different modes from OMSSA’s p-values.

**Figure 9:**
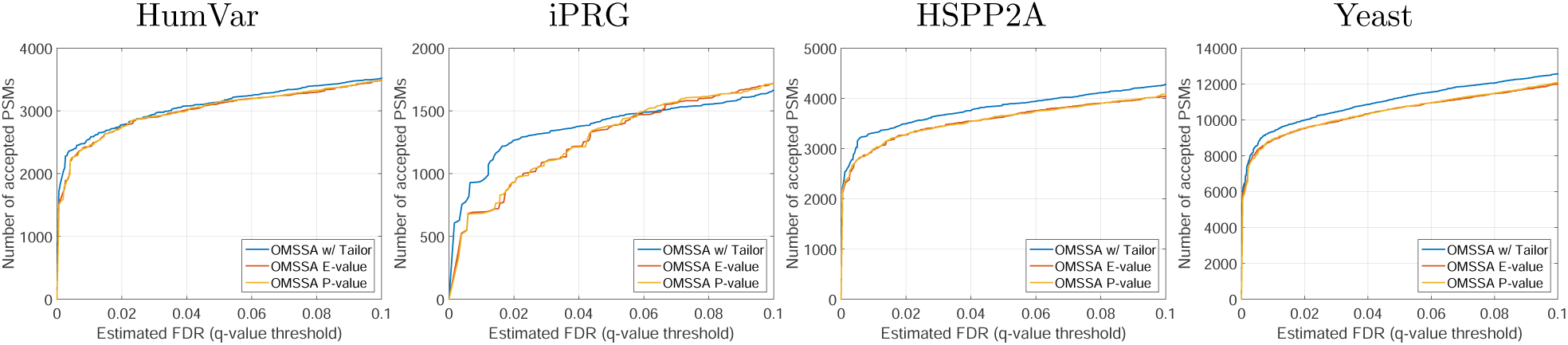
Spectrum annotation results obtained with OMSSA program. Each plot corresponds to different datasets indicated on top.

We note that we did not re-calibrate the OMSSA’s E-values with the Tailor method because OMSSA’s p-values and E-values are highly correlated and would result in the same outcome.

### Combination with post-processing methods

The Tailor method, similarly to any other score calibration methods, can be combined with any postprocessing methods such as Percolator^33^ and PeptideProphet^40^ in order to obtain additional spectrum annotations at given FDR. We tested Percolator with calibrated scores and displayed the results on Figure 10. We found that combining digestion- and charge-state-specific information with scores calibrated with XPV and Tailor using Percolator can result in additional number of PSMs over the entire 0-10% FDR range. Overall, the performance of the Tailor score calibration with Percolator post-processing remains on par with exact calibration methods with Percolator and outperforms other methods such as OMSSA, X!Tandem, and MS-GF+ on our benchmark, shown on Figure 10. Table 3 shows the number of annotations obtained at 1% FDR level and the relative improvement obtained by the application of Tailor compared to other methods.

**Table 3:**
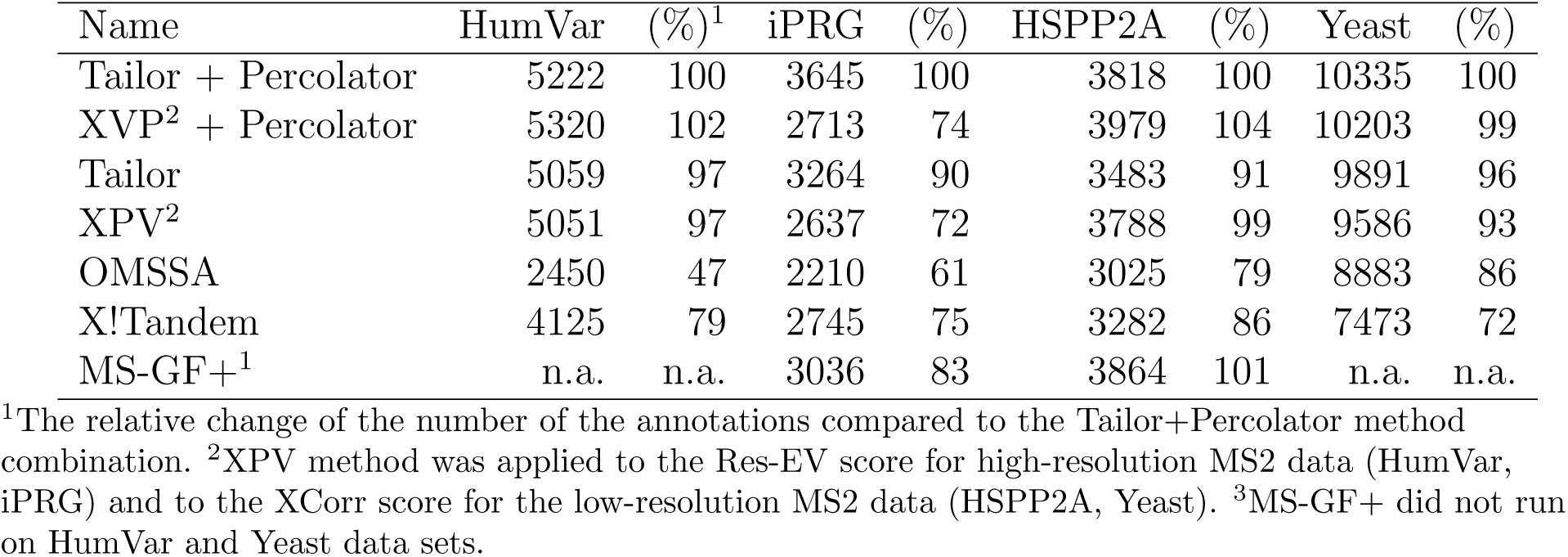
Number of spectrum annotations at FDR level of 1%.

**Figure 10:**
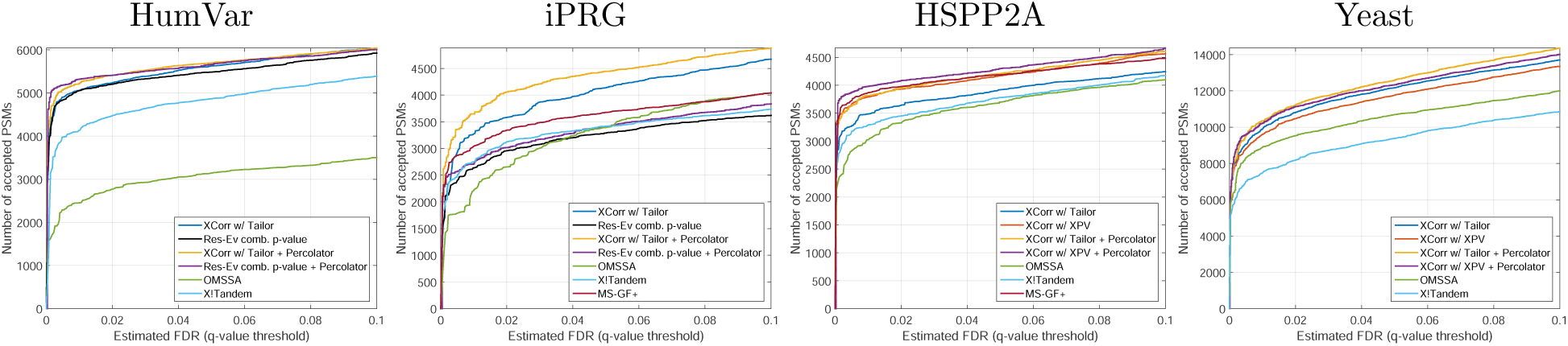
Overall results. Each plot corresponds to different datasets indicated on top. Note that MS-GF+ did not run on HumVar and Yeast data sets.

## Conclusions

In this article, we introduced a new, heuristic score calibration method, called Tailor, for score functions employed in database searching mass spectrometry. The Tailor method calibrates the PSM scores relatively to the tail of the empirical null distribution of any experimental spectrum. The Tailor method does not require additional computational steps, can be performed based on the observed PSM scores, and its performance is on par with the computationally expensive exact methods; however, Tailor runs around 100 times faster and could save substantial amount of CPU time and energy. We hope that, its simplicity and robustness will make the Tailor method a standard option in many database searching software programs. The Tailor method is freely available in the open source Crux mass spectrometry toolkit (http://crux.ms).

## Acknowledgments

We thank William Stafford Noble and the Crux development team for hosting Tailor method in the Crux mass spectrometry toolkit at http://crux.ms.

## Author contributions

A.K.F. designed the experiments, performed data analysis, and wrote the manuscript. P.S. developed scripts and programs and performed data analysis. All authors read and approved the final manuscript.

## Competing financial interests

The authors declare no competing financial interests.

## Supporting Information Available

- Supplementary Note S1: additional information to Percolator search.
- Supplementary Figure S1. Spectrum identification with Percolator and Tide in decoy-decoy search scenario.
- Supplementary Note S2: Information for classification of target-decoy theoretical spectra.
- Supplementary Figure S2: Model architectures of the logistic regression and the vanilla neural network models.
- Supplementary Figure S3: ROC results on the classification of target against reversed decoy peptides.
- Supplementary Figure S4: ROC results on the classification of target against shuffled decoy peptides.
- Supplementary Figure S5: ROC results on the classification of target-against-target and decoy-against-decoy peptides.
- Supplementary Note S3: Information for training weighted XCorr score function with logistic regression.
- Supplementary Note S4: Information about the *Plasmodium falciparum* data.
- Supplementary Note S5: Notes and details on the Target-decoy peptide correlation using synthetic data.
- Supplementary Table S1:Statistics of the peptide sets using various peptide generation methods.
- Supplementary Figure S6: Correlation between target and decot peptides obtained with various peptide generation methods.
- Supplementary Note S6: Notes on *p*−value validation.
- Supplementary Note S7: Notes on the correction for protein-level shuffling.
- Supplementary Figure S7: ROC analyses of protein-level shuffling.

