## Supplementary Material for "Tailor: non-parametric and rapid score calibration method for database search-based peptide identification in shotgun proteomics"

**Supplementary Note S1: Database search program parameters**

**Tide-search.** The tide-search program [1] from Crux [2] version 3.2.8aa66a2 was downloaded and used in Linux to execute XCorr and Res-ev score functions and their corresponding exact p-value (XPV) calculation methods. We note that the raw XCorr score function and the XCorr score function for the XPV algorithms are carried out with different code, and they are also slightly different. The raw XCorr considers only singly charged fragment ions for precursors of charge 1+ or 2+; however, it considers singly and doubly charged fragment ions for a precursor with a charge state larger than 2+ [1]. In contrast, the XCorr score function in the XVP calculation considers the fragment ions of charge $n-1$ for a precursor ion if charge $n>1$ [3]. This discrepancy can result in different discrimination power for the score functions making the comparison of their calibration power more difficult. Therefore, we adjusted the source code of the XPV method to raw XCorr score function so that they become equivalent. A patch file performing this code update is available in the supplementary materials.

The tide-search program was run with the following parameters: `min-peaks=10, use- neutral- loss-peaks=F, concat=T, max-precursor-charge=9`. The `precursor-window, precursor- window- type`, and `mz-bin-width` were specified with respect to the input spectrum dataset. The p-values for the XCorr scores were obtained using the `exact-p-value=T` parameter, and the p-values obtained were adjusted to the multiple hypothesis testing using the Šidák correction [4]. The Res-ev score function [5] was used for high-resolution MS2 data (HumVar, Malaria, iPRG).

Tide-index generated the preprocessed peptide index files with the following parameters: `decoy-format= none`, `enzyme= custom-enzyme, custom-enzyme=[Z]|{Z}` and `missed-cleavages=0`. Modifications were specified for the given taxon. All other parameters were used with their default values.

**X!Tandem.** The X!Tandem version of 2015 was used with the following parameter settings: `cleavage site=[Z]|{Z}, refine=No, minimum peaks=10, total peaks=50, output,results=all`. Note that, the X!Tandem defines the fragment ion peak match tolerance as $|p-m|<t$; therefore, `mz-bin-width=1.0005079` would be equivalent to `fragment monoisotopic mass error=0.5 Da`, which leads to roughly the same spectrum discretization step. For high-resolution data, `fragment monoisotopic mass error=0.01 Da` was specified, which is equivalent to `mz-bin-width=0.02`. Other dataset-specific parameters were specified accordingly. The X!Tandem program uses the HyperScore scoring function defined as $\text{HyperScore}(s,t) = N_b!N_y!M(s,t)$, where $N_b!$ and $N_y!$ denotes the factorial of the number of matching $b$ and $y$ ions in the theoretical spectrum.

The Tailor score calibration of the HyperScore scores was executed on the output of the X!Tandem program (.xml file), and it was implemented as a Python script. We note that the experimental spectra, that did not have at least 30 candidate peptides were excluded from our analysis. We also note that X!Tandem

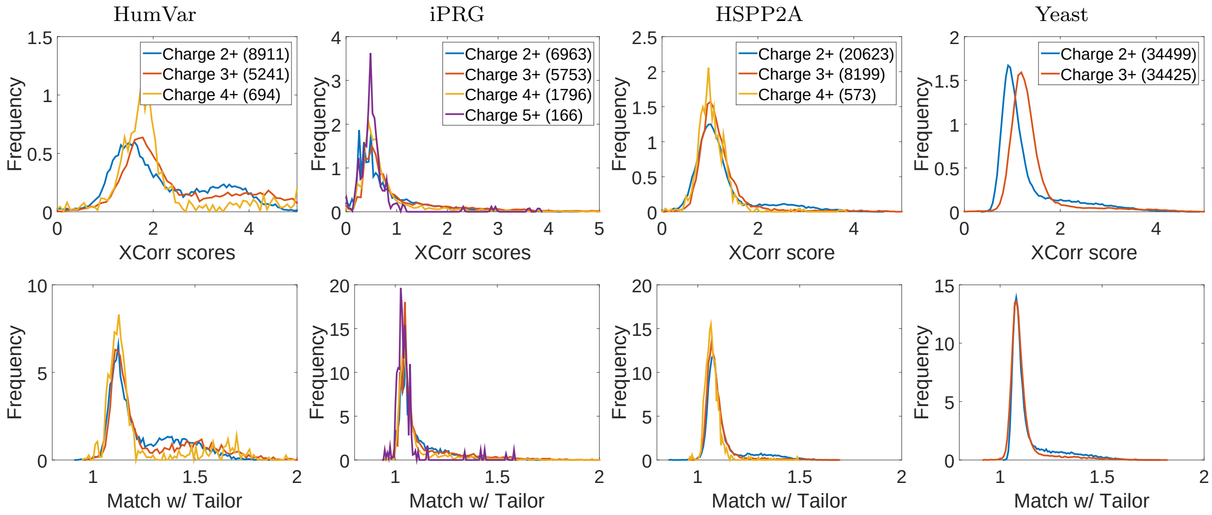


Figure S1: Score distributions of the XCorr (top row) and the Tailor (bottom row) methods with respect to precursor ion charge states. The columns correspond to different datasets and share a figure legend. The distributions were smoothed with the Gaussian kernel.

does not report the PSMs of the candidate peptides individually for every experimental spectrum, it reports only the histogram of the empirical cumulative survival distribution of the HyperScore scores, and all relevant information needed for Tailor, such as the number of the candidate peptides and the raw HyperScore scores, was extracted from this histogram.

**OMSSA.** The OMSSA program version 2.1.9 was downloaded from `ftp://ftp.ncbi.nlm.nih.gov/pub/lewisg/omssa/2.1.9/`. For this program, the fasta files were preprocessed with the `makeblastdb` program of the BLAST toolbox version 2.9.0+ with the parameter `-dbtype prot`. The omssacl program was run with the following parameters: `-i 1,4` ($b$ and $y$ ions used), `-mnm` (disabling protein N-terminal methionine cleavage), `-mf 3` (static carbamidomethylation of cysteine), `-e 11` (disabling in silico digestion) `-he 99999999` (maximum e-value), `-hl 9999` (retain top 9999 hits), `-zcc 1` (believe precursor charge), `-tem 0 -tom 0` (monoisotopic precursor and fragment ions), `-tez 1` (linear charge dependency of precursor charge), texttt-zh 9 (max charge state), `-v 0` (number of missed cleavages 0), and `-oc` (csv output). The precursor ion (`-te, -teppm`) and the fragment ion tolerance (`to`) parameters were specified with respect to the input spectrum dataset. All other parameters remained at their default value.

We note that we did not implement Tailor inside the OMSSA program, but as a python script, which post-processes the comma-separated `csv` output of OMSSA. OMSSA was parameterized to report all PSMs corresponding to all candidate peptides for all experimental spectra so that the Tailor method could extract the information needed. We also note that the experimental spectra that have fewer than 30 candidate peptides were excluded from our analysis.

### Supplementary Note S2: Supplementary figures

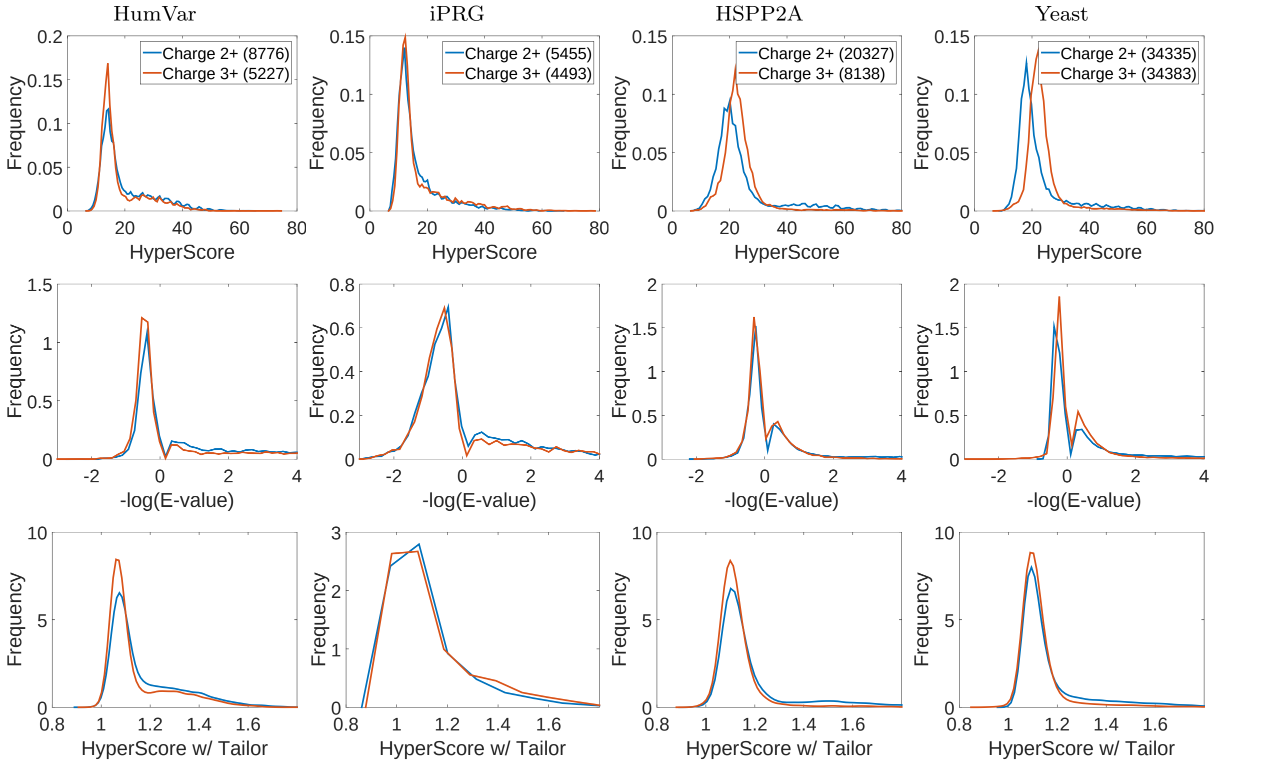


Figure S2: Score distributions of the HyperScore (top row), log E-value (middle), and the HyperScore calibrated with Tailor (bottom row) methods with respect to precursor ion charge states. The columns correspond to different datasets and share figure legend. The distributions were smoothed with the Gaussian kernel.

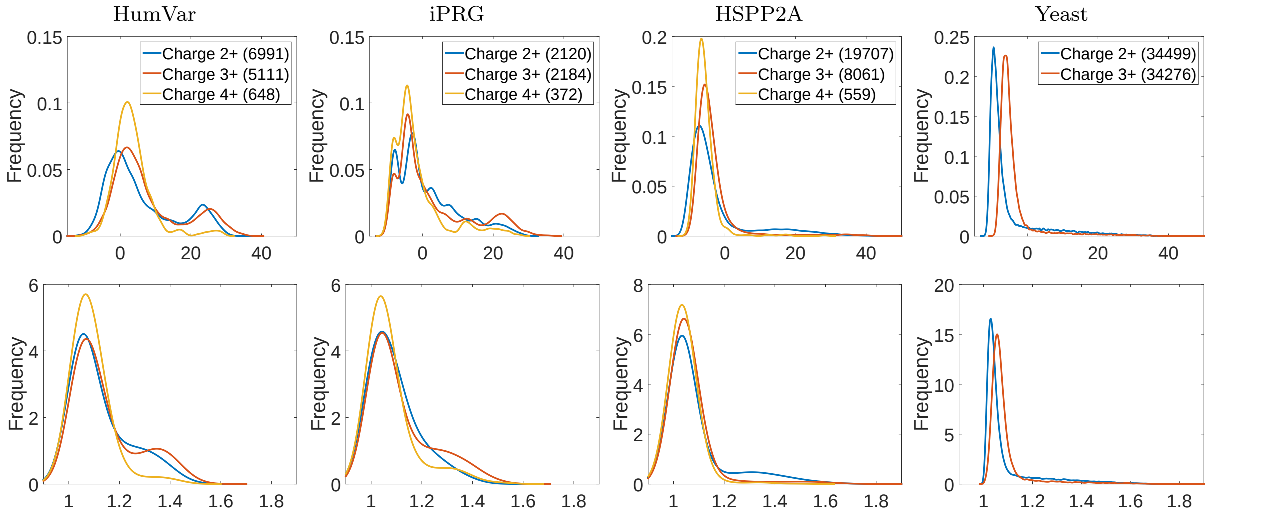


Figure S3: Score distributions of the top-scoring PSMs with respect to the charge state of the precursor ions. Each column corresponds to a dataset, plots in the top row correspond to OMSSA's p-value, and plots in the bottom row corresponds to the Tailor method. The distributions were obtained with kernel smoothing using the Gaussian smoother.
